# CyStainer: A transformer-based variational autoencoder for robust marker imputation in high-parameter cytometry

**DOI:** 10.64898/2026.06.30.735235

**Authors:** Konstantin Ivanov, Mouhamad Al Moussawy, Frederik Kirk, Rounioja Samuli, Olli Lohi, Lars Rønn Olsen, Signe Modvig, Ville Hautamäki, Merja Heinäniemi

**Affiliations:** School of Medicine, University of Eastern Finland, Kuopio, Finland; Department of Clinical Immunology, Copenhagen University Hospital Rigshospitalet, Copenhagen, Denmark; Department of Clinical Medicine, Copenhagen University, Copenhagen, Denmark; Department of Hematology, Fimlab Laboratories, Tampere, Finland; Faculty of Medicine and Health Technology, Tampere University Hospital, Tampere, Finland; School of Computing, University of Eastern Finland, Joensuu, Finland

**Keywords:** cytometry, artificial intelligence, marker imputation

## Abstract

High parameter cytometry is essential for clinical diagnostics through precise immune cell profiling, improved patient stratification, and monitoring, while also enhancing the understanding of cellular responses in disease and therapeutic contexts. The amount of cytometry data is growing fast, and with that, the need to merge different datasets for unified analysis. Here, we present CyStainer, a transformer-based variational autoencoder that demonstrates competitive or superior performance to existing methods on several key tasks related to marker prediction. As a key novelty, we demonstrate that CyStainer can impute markers without having a set of shared backbone markers. We performed several benchmarks using real-world FACS, CyTOF, InfinityFlow and CITE-seq datasets to show that CyStainer is a robust and flexible tool for panel merging, marker imputation, dataset integration and virtual staining of unseen samples.

**Graphical Abstract:** 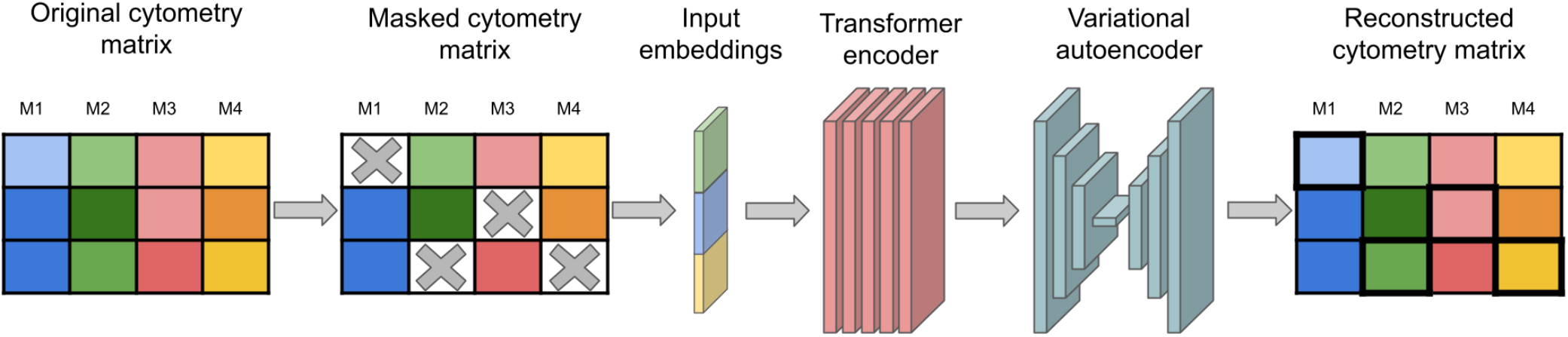

## Introduction

High parameter cytometry is an advanced approach for high dimensional analysis of heterogeneous cell populations at cellular resolution. It allows measurement of a wide range of targets located on the cell surface, in the cytoplasm, or within the nucleus, typically proteins but also other molecules such as carbohydrates and RNA (McKinnon, 2018; Sun et al., 2020). By using antibodies conjugated to distinct reporter molecules (fluorophores) for conventional and spectral flow cytometry (fluorescence-activated cell sorting, FACS), or heavy metal isotopes for mass cytometry (cytometry by time-of-flight, CyTOF) these cytometry technologies enable the simultaneous quantification of up to 50-60 cellular features. Modern instruments can analyse millions of cells in a short time with relatively simple sample preparation. Sequence-tagged antibodies extend these approaches to multimodal characterisation, including methods such as Cellular Indexing of Transcriptomes and Epitopes by sequencing (CITE-seq) to measure the whole transcriptome and several hundred proteins simultaneously. The broad adoption of these methods generates vast amounts of data, creating opportunities for comprehensive big data analyses.

Since its introduction in the 1970s, flow cytometry technology has become widely used in both research and clinical applications, particularly in hematology, where the nature of the samples facilitates analysis of individual cells. Today, high parameter cytometry is an essential part of routine clinical practice. For example, flow cytometry is indispensable for detecting minimal residual disease in hematologic malignancies, enabling risk estimation and assessment of the treatment response (Van Dongen et al., 2012; Li, 2022). However, the number of markers that can be measured in a single analysis remains limited, especially in the clinical setting, and instead several panels are often run in parallel from the same sample. Typically, the individual panels range from 6-15 detectable markers, requiring imputation for capturing the full marker profile. Moreover, since panel design is often highly diagnosis specific, a large number of unique marker panels exist that may vary by time point or measurement site, complicating dataset integration and analysis.

A combination of experimental and computational approaches to address the above technical and practical limitations of cytometry exist, such as the Infinity Flow method, where a specific experimental setup is used to incrementally collect information across hundreds of markers in combination with gradient boosting regressors (GBR) to infer missing markers (Becht et al., 2021). Marker imputation methods that require no special experiment design and are directly applicable to existing data include statistics and artificial intelligence (AI) based approaches that utilise partially overlapping markers across measurements. In a setup where panel consistency is preserved, a masked autoencoder (cyMAE) approach was shown to enhance regression imputation and to enable prediction of markers in unseen data (Kim et al., 2024). Other approaches that can mitigate technical batch effects and combine data across platforms include the cyCombine method which performs integration followed by a probability-based imputation at cluster level and cytoVI that represents a recent neural network approach using a variational autoencoder (VAE) architecture (Pedersen et al., 2022; Ingelfinger et al., 2025).

Despite these developments, two fundamental challenges remain unsolved: the existing solutions that allow predicting unseen data require a fixed configuration of backbone markers while more flexible integration methods involve multistep iterative integration pipelines. This poses a severe limitation to what new datasets the models can be directly applied to without re-training, which limits feasibility in the big data context.

In this study, we propose CyStainer, a deep learning framework that is compatible with several cytometry platforms and capable of handling cross-study and cross-panel variability while maintaining strong imputation accuracy. By utilising a transformer encoder CyStainer can capture the complex, nonlinear relationships and co-expression patterns between markers. Importantly, this architecture accommodates training across panels in the absence of shared backbone markers. In combination with a VAE, the model learns a robust, compressed latent representation of cell states, aiding generalisation and reducing overfitting. These improvements allow CyStainer to achieve state-of-the-art accuracy across multiple cytometry platforms. We provide a comprehensive assessment of model performance based on cell-wise imputation accuracy, distributional accuracy, conditional distributional accuracy within pre-defined groups of cells and biological accuracy. CyStainer shows superior performance in preserving the biological interpretation and generalisation when backbone markers are missing.

## Methods

### Datasets

#### Dataset selection and Preprocessing

The datasets selected for the initial model testing include two CyTOF datasets, CytoBackBone and COVID-19, used previously in the development and testing of the cyCombine and cyMAE methods, respectively. The CytoBackBone healthy whole blood dataset was acquired from Flow Repository (FR-FCM-ZYV2). The COVID-19 blood datasets from the cyMAE study were acquired from Pennsieve Platform for Data Management. Both datasets contain filtered data and only the arcsinh transformation with a factor of 5 was used as a pre-processing step. Secondly, we chose a dataset representing the clinical cytometry context from Flow repository and retrieved the acute lymphoblastic leukemia (ALL) bone marrow flow cytometry dataset (FR-FCM-Z68U) that represents diagnostic profiles collected across several different hospitals. This data was already filtered and the same arcsinh transformation with factor 100, followed by a min-max scaling, were applied to preprocess the data. In addition, saturated values, e.g. equal to 0 or 1, were removed. Thirdly, to test imputation performance for a large set of markers, we retrieved the Infinity Flow bone marrow dataset from Zenodo (15065910). Negative scatter values were removed, arcsinh transformation with a factor of 500 was applied, followed by min-max scaling and removal of saturation at 0 and 1 as for the clinical flow cytometry dataset. Fourth, the NeurIPS CITE-seq dataset was downloaded from the Gene Expression Omnibus data repository (GSE194122). This dataset contains a high number of markers, samples across several donors that were processed at different laboratories (sites 1-4) and pre-annotated cell types, making it well-suited for testing model generalization and preserving biological interpretation (Luecken et al., 2021). The CITE-seq data was already filtered and log-normalized. Therefore, only min-max scaling was applied in preprocessing.

#### Experimental setups for model testing

From the CytoBackBone dataset the panels A, B, C and D were used for training of the CyStainer model and panel E was used for the leave-one-out test where each marker in turn was dropped and predicted. For comparison, CyCombine results were generated with all panels with one marker dropped from panel E in turn which is not equivalent (more data used in training) but the closest setup that this method supported. For testing CyStainer in the backbone marker prediction task, we used the same panels for training and calculated performance metrics for the entire subset of panel E markers that did not overlap between all panels. From the additional external clinical dataset (ALL), the data from one hospital (Niñ o Jesú s Hospital, 96 samples) was used for training and the remaining 20 samples for a leave-one-out test. In COVID-19 blood CyTOF datasets we reproduced the original training and test scheme used by the cyMAE developers: The “acute2020” datasets were used for training while the “vaccine” and the “acute2021” datasets were used for testing with the task to predict the held-out T cell markers (CD27, CD28, CD45RA, CD45RO, CD127,CD197, TCRgd). The Infinity Flow dataset was divided into training dataset (7 donors) with presence of backbone and unique markers and a testing dataset (2 donors) with only backbone markers. In the NeurIPS dataset we tested the model performance in the presence of technical batch effects. To achieve this, one group of markers was dropped from each site: from site 1 - T cell markers; from site 2 - B cell markers; from site 3 - natural killer (NK) and innate lymphoid cell (ILC) markers; from site 4 - monocyte, macrophage and dendritic cell markers. The list of markers for each group is available in Supplementary Table S1. Furthermore, B cell annotations in the NeurIPS dataset were simplified prior to evaluation. Subpopulations differentiated solely by immunoglobulin kappa constant (IGKC) expression (e.g., IGKC+ and IGKC-) were merged into their respective parent cell types (Naive CD20+ B, B1 B, Plasmablast, and Plasma cell). We specifically selected cytoVI as the baseline for this benchmark because, in the presence of such complex technical batch effects, only CyStainer and cytoVI possess the architectural flexibility to perform targeted marker imputation across distinct batches. Consequently, both models were trained on all sites, and the omitted markers were subsequently predicted for each respective missing site.

To test adaptation to a new unseen batch we dropped the same marker groups as before from the sites 1, 2 and 3 and trained the model without site 4. After that, the CyStainer model was fine-tuned with site 4, where each of the three groups of markers that were omitted in turn during the training were predicted for site 4.

### Model architecture

The CyStainer architecture, as depicted in Fig. 1, has features embedders for the numerical cytometry values, categorical marker identities, and batch values. First, the expression values for n markers are projected to a hidden layer via a fully connected layer without activation functions and a normalisation layer. Then batch and marker identities are embedded using a standard embedding layer available in PyTorch. These three embeddings are concatenated to form the combined input representation that is fed into the transformer encoder. The core of the transformer encoder is the attention mechanism that enables learning complex relationships between markers (Vaswani et al., 2017):

**Figure 1.**
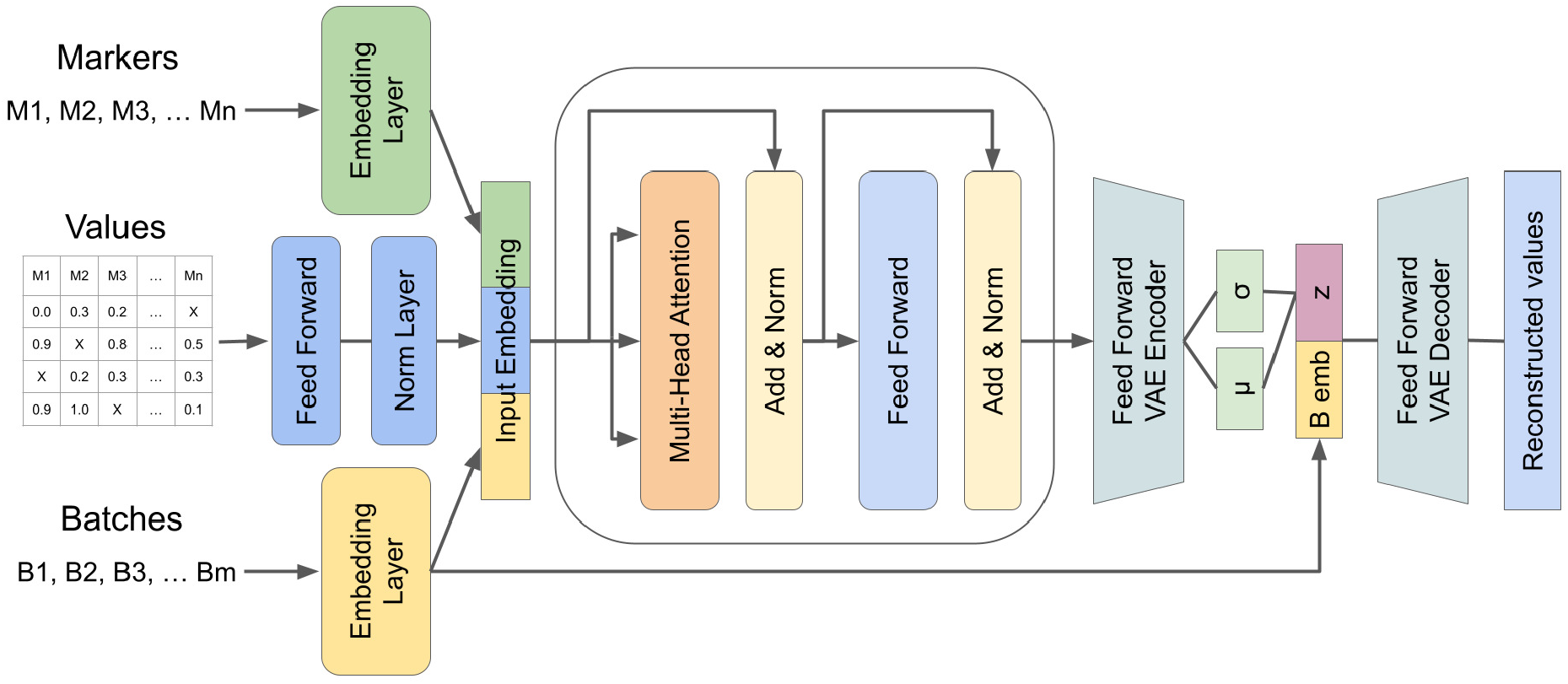
Overview of the CyStainer architecture. Categorical inputs (marker and batch identities) and numerical expression values are embedded and processed through a Transformer encoder utilizing multi-head attention to capture non-linear marker relationships. The output is projected into a latent space via a VAE, allowing for robust batch correction and reconstruction of the full cytometry matrix.

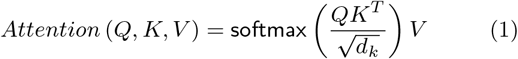

The attention mechanism was implemented using multi-head attention:

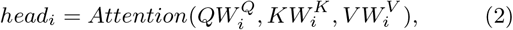

where the matrix *Q* contains queries, matrices *K,V* contain key-value pairs, 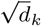 represents key dimension and serves as a scaling factor and *W* represents weights. The transformer encoder output is flattened and projected into a latent space by the VAE encoder, aiding in generalization and allowing batch correction (Kingma and Welling, 2013). The encoder consists of two fully-connected layers followed by a dropout layer where a SiLU activation function is used in each (Hendrycks and Gimpel, 2016; Householder, 1941). Finally, a sample drawn from the latent distribution is passed to the VAE decoder to reconstruct the full expression panel. The decoder consists of three fully-connected layers first two layers are followed by a dropout layer and SiLU activation. The model was implemented using PyTorch.

### Masking strategy

During training, we employ a separated masking function to independently regularize shared and unique markers. Let *M*_shared_ and *M*_unique_ represent the sets of valid, unpadded markers in a given sequence. We define maximum masking rates *r*_shared_ and *r*_unique_ for these sets (by default 0.3 and 1.0, respectively). For each sequence in a batch, the maximum number of maskable tokens is calculated as *N*_shared_ *× r*_shared_ and *N*_unique_ *× r*_unique_. The actual number of tokens to mask for each group is then randomly sampled from a uniform distribution bounded by [0, *N*_max_], where *N*_max_ is the respective calculated maximum for that group.

### Training

Huber loss was employed as the regression loss function, calculated per data point as:

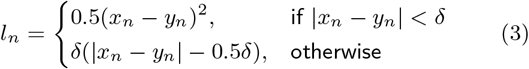

and averaged over the entire batch:

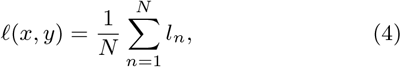

where *N* is the total number of samples in the batch, *x*_*n*_ is the *n*-th element of the model prediction, *y*_*n*_ is the *n*-th element of the ground truth, and *δ* is a hyperparameter defining the threshold between the quadratic and linear penalties.

To ensure that the latent distribution is close to a standard normal distribution we used the Kullback–Leibler divergence as a second loss function with a default weight of 0.0001. First, we define the prior over the latent variables as a standard isotropic Gaussian, *p*(**z**) = *N* (**0, I**). The approximate posterior is modeled as a diagonal Gaussian, *q*(**z**|**x**) = *N* (***µ***, diag(***δ***^2^)). The KL divergence is averaged over the batch of size *N* and summed across the latent dimensions *D*, yielding the closed-form objective:

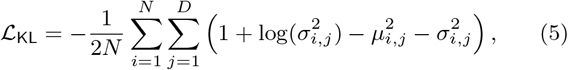

where *N* is the total number of samples in the batch, *D* is the latent dimension, *µ*_*i,j*_ is the mean, *σ* _*i,j*_ is the variance and *N* (**0, I**) is the prior. The CyStainer model was trained until loss function convergence. We used AdamW as optimizer (Loshchilov and Hutter, 2017), with a learning rate of 10^*−*4^ for initial training and 10^*−*3^ for fine-tuning. During training, we set a maximum of 300 epochs and employed early stopping with a loss tolerance of 10^*−*4^ and a patience of 3 epochs. For the fine-tuning phase, the maximum number of epochs was reduced to 50, while maintaining the same early stopping scheme. To ensure convergence while training CyStainer on the NeurIPS dataset, the loss tolerance was adjusted to 10^*−*6^ with a patience of 5 epochs.

### Fine-tuning and marker imputation

To apply a pre-trained CyStainer model to new, unseen datasets subject to technical variation or distinct batch effects, we implemented a targeted fine-tuning strategy. When new samples are introduced, the model dynamically expands its batch vocabulary by appending additional indices to the existing batch embedding layer while strictly preserving the previously learned weights.

During the fine-tuning phase, the core network architecture elements, including the transformer encoder, the VAE blocks, the value embeddings, and the marker embeddings, are frozen. This ensures that the robust, global marker co-expression patterns learned during initial training are retained. Gradient updates are restricted only to the expanded batch embedding layer. In this manner, the model e”ciently learns the specific technical variations of the new batches without the computational overhead or the risk of catastrophic forgetting associated with full model retraining.

Marker imputation is performed by passing the masked or partially observed marker profiles through the network, projecting them into the latent space, and decoding them to reconstruct the complete expression panel. Furthermore, CyStainer facilitates explicit batch correction during this inference step. To align cells across different distributions, the observed data is first encoded using its original source batch index to extract the latent mean representation. This representation is subsequently passed to the decoder alongside a user-designated target reference batch index, effectively translating the imputed marker values into the target batch distribution.

### Performance metrics

#### Cell-wise accuracy

To evaluate cell-wise accuracy of the models we used two metrics: Pearson correlation and R-squared. Pearson correlation metric function available from the scipy python package is calculated as follows:

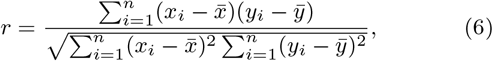

where *r* represents the Pearson correlation coefficient, *n* represents the total number of samples, *x*_*i*_, *y*_*i*_ contain the individual sample values such as predicted and true values, and 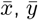 represent the mean values of *x* and *y* respectively. R-squared metric function available from the scikit-learn python package is calculated as follows:

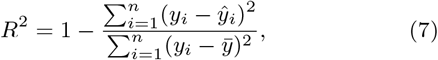

where *R*^2^ represents the coefficient of determination (R2 score), *n* represents the total number of samples, *y*_*i*_ and *Ŷ*_*i*_ contain the individual sample values for the true and predicted values respectively, and 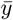 represents the mean of the true values *y*.

#### Distributional accuracy

To evaluate distributional accuracy we used the Earth Mover Distance (EMD) metric function, specifically the Wasserstein distance available from the scipy python package calculated as follows:

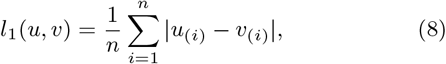

where: *l*_1_(*u, v*) represents the first Wasserstein distance. *n* represents the total number of samples in each array. *u*_(*i*)_ represents the *i*-th value of array *u* after it has been sorted in ascending order. *v*_(*i*)_ represents the *i*-th value of array *v* after it has been sorted in ascending order.

#### Conditional distributional accuracy

To test whether the marker distribution was preserved for a set of similar cells (conditional accuracy), we performed clustering using the Leiden algorithm on the backbone markers (Infinity Flow dataset). Clusters 17, 18, and 19 were omitted from this analysis because their cell counts were exceptionally low. The metrics were calculated between predicted and real values within each cluster and summarized for the top 5 markers defined for each cluster based on the Wilcoxon test.

#### Biological accuracy

To assess whether the imputed data preserves the separation of ground-truth cell types to a similar level as the original measured marker profile, we used the Neurips data with available cell type labelling from the original study. The original and imputed profiles were clustered using the Leiden algorithm with resolution starting with 0.01 and then ranging from 0.1 to 1.5 with step 0.1. As performance metrics, we used the adjusted rand index (ARI) and normalized mutual information (NMI) metrics available in scikit-learn. The ARI is calculated as follows:

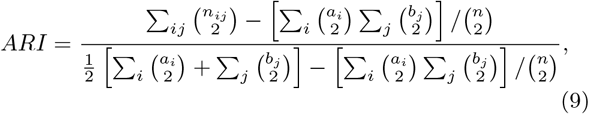

where *n* represents the total number of samples, *n*_*ij*_ represents the number of samples present in both the true class *i* and predicted cluster *j, a*_*i*_ is the total number of samples in true class *i*, and *b*_*j*_ is the total number of samples in predicted cluster *j*. The NMI is calculated as follows:

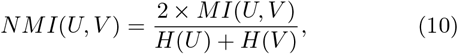

where *U* represents the ground-truth cell type labels, *V* represents the predicted cluster labels, *MI*(*U, V*) is the mutual information between the two label assignments, and *H*(*U*) and *H*(*V*) represent the entropy of the respective assignments.

## Results

### Comparison results on benchmark datasets

As a first comparison against existing tools, we retrieved the whole blood CytoBackBone CyTOF dataset which was originally used for evaluating the cyCombine method (Pereira et al., 2019). The partially overlapping panels A, B, C and D were used in training and panel E that includes all markers was used as a test to evaluate model performance for each marker inturn. The performance in the leave-one-out test benchmarked against cyCombine is shown in Figure 2 and Table 1a. The new CyStainer model demonstrates superior performance in cell-wise imputation accuracy (r 0.500 vs 0.319 and R-squared 0.246 vs -0.365, scatter plots Fig. S1), while having slightly worse scores for the distributional accuracy assessed using EMD (0.217 vs 0.089). To demonstrate that CyStainer’s performance translates to routine clinical flow cytometry, we repeated the experiment in real-world clinical flow cytometry data from acute lymphoblastic leukemia (ALL) (Table 1b and Fig. 3) obtaining consistent differences between the two methods (Chulián et al., 2023).

**Table 1.** Comparison of imputation across multiple datasets. Values represent mean metrics and standard deviations across tested markers.

| (a) Healthy whole blood CyTOF leave-one-out panel merging |  |  |  |
| --- | --- | --- | --- |
| | $r$ | $R^2$ | EMD |
| CyStainer | **0.500**±0.175 | **0.246**±0.184 | 0.217±0.094 |
| cyCombine | 0.319±0.193 | -0.365±0.377 | **0.089**±0.023 |

| (b) Healthy whole blood CyTOF panel merging without backbone markers |  |  |  |
| --- | --- | --- | --- |
| | $r$ | $R^2$ | EMD |
| CyStainer | **0.522**±0.165 | **0.233**±0.245 | 0.196±0.068 |
| cyCombine* | 0.291±0.154 | -0.463±0.328 | **0.110**±0.046 |

\*For reference, performance in leave-one-out marker imputation is shown for cyCombine.

| (c) ALL flow cytometry leave-one-out panel merging |  |  |  |
| --- | --- | --- | --- |
| | $r$ | $R^2$ | EMD |
| CyStainer | **0.782**±0.124 | **0.590**±0.214 | 0.036±0.017 |
| cyCombine | 0.554±0.123 | 0.114±0.242 | **0.021**±0.013 |

| (d) Healthy bone marrow Infinity Flow panel merging |  |  |  |
| --- | --- | --- | --- |
| | $r$ | $R^2$ | EMD |
| CyStainer | **0.664**±0.149 | **0.360**±0.285 | 0.063±0.025 |
| cyCombine | 0.381±0.164 | -0.403±0.710 | **0.046**±0.025 |
| cytoVI | **0.664**±0.176 | 0.343±0.298 | 0.075±0.027 |
| GBR | 0.637±0.173 | 0.306±0.339 | 0.065±0.027 |

| (e) Healthy bone marrow CITE-seq panel merging |  |  |  |
| --- | --- | --- | --- |
| T cell markers |  |  |  |
| | $r$ | $R^2$ | EMD |
| CyStainer | **0.743**±0.235 | **0.484**±0.373 | **0.054**±0.029 |
| cytoVI | 0.735±0.243 | 0.398±0.379 | 0.075±0.028 |
| B cell markers |  |  |  |
| | $r$ | $R^2$ | EMD |
| CyStainer | **0.684**±0.156 | -0.480±2.644 | **0.053**±0.036 |
| cytoVI | 0.675±0.161 | **-0.368**±2.461 | 0.059±0.042 |
| NK and ILC cell markers |  |  |  |
| | $r$ | $R^2$ | EMD |
| CyStainer | **0.475**±0.164 | **-0.054**±0.372 | **0.073**±0.033 |
| cytoVI | 0.467±0.140 | -0.079±0.342 | 0.079±0.031 |
| Monocytes, macrophages and DC cell markers |  |  |  |
| | $r$ | $R^2$ | EMD |
| CyStainer | 0.734±0.170 | **-0.554**±1.977 | **0.053**±0.029 |
| cytoVI | **0.737**±0.171 | -0.735±2.149 | 0.056±0.039 |

**Figure 2.**
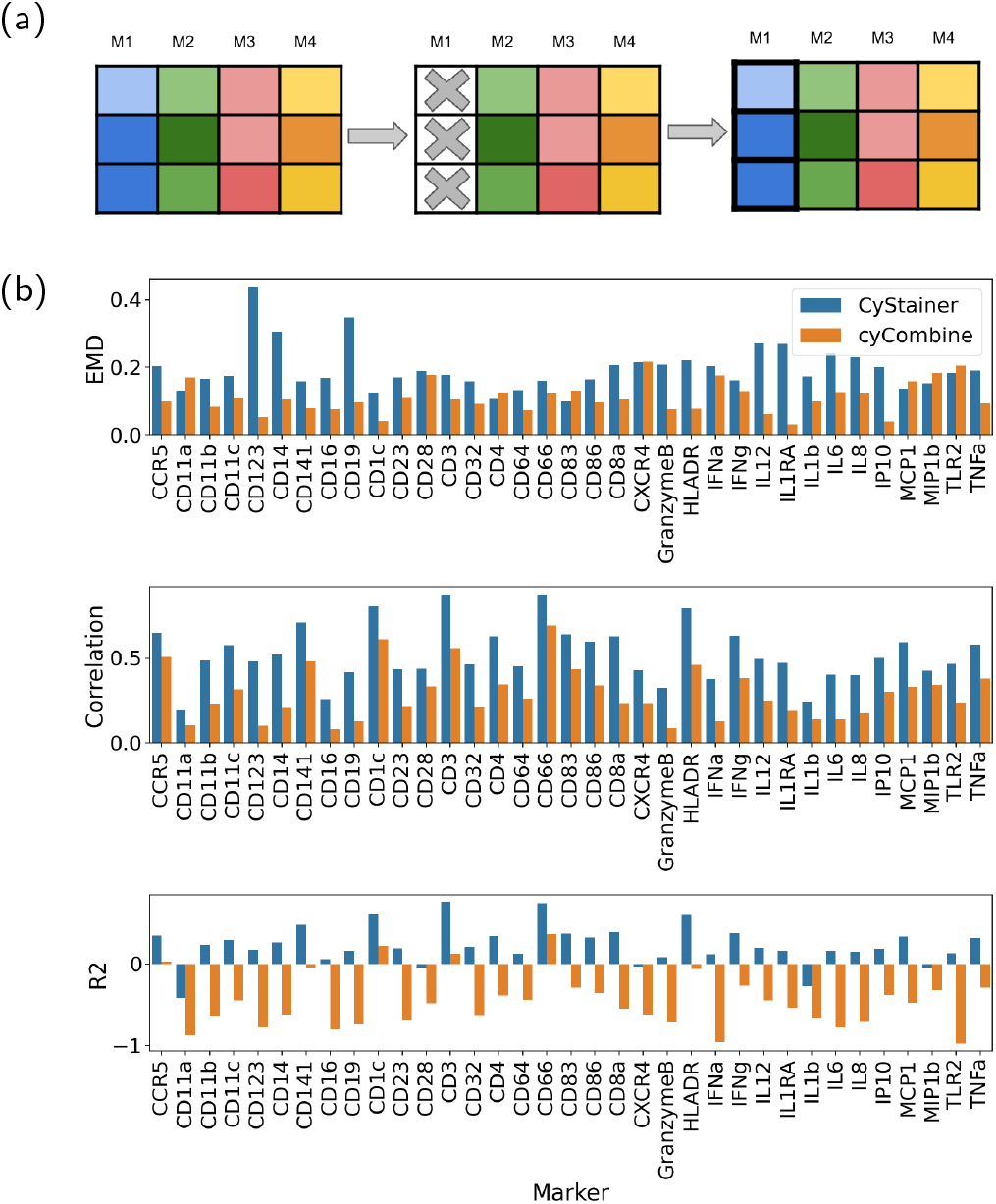
Performance comparison of CyStainer and cyCombine in leave-one-out marker imputation using panel E. (a) Leave-one-out scheme. (b) EMD, Pearson correlation and R-squared metrics marker wise.

**Figure 3.**
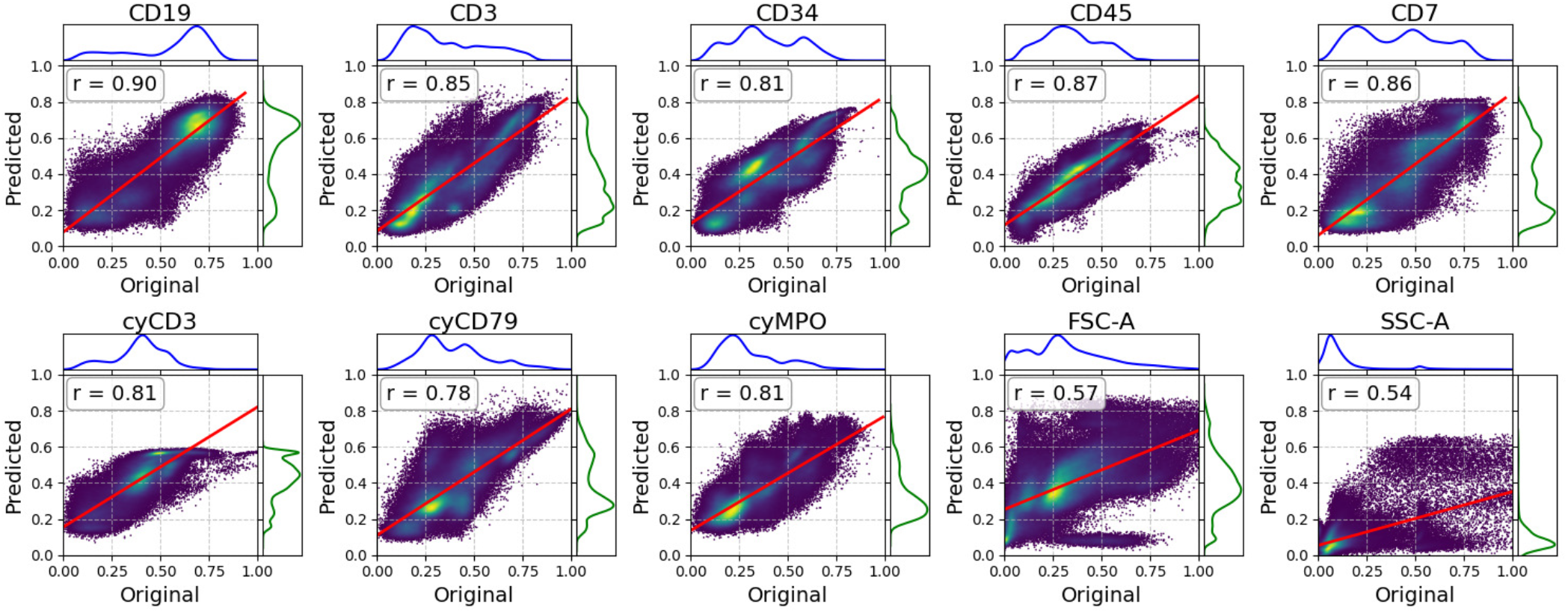
Regplots of real versus CyStainer predicted values with Pearson correlation values for each marker in leave-one-out test for ALL flow cytometry dataset

In the previous benchmark, the model training and test data originated from the same study. To demonstrate CyStainer performance in an out-of-distribution prediction task, we chose cytometry datasets which were originally used in the study by Kim et al. (2024) to benchmark the cyMAE model. We evaluated CyStainer against the combination of the cyMAE and GBR proposed by the authors (cyCombine is incompatible with this experimental setup). The study has three CyTOF collections of blood samples: two of which are from acute COVID-19 patients from 2020 and 2021, and the third one is the vaccinated with COVID-19 vaccine healthy donors. Previously, cyMAE was shown to improve imputation quality with the help of GBR compared to regressor alone. We reproduced the benchmarking test where training is done using the “acute2020” CyTOF dataset from COVID-19 patients and prediction of T cell markers is evaluated on the “acute2021” and “vaccine” datasets. The target markers CD27, CD28, CD45RA, CD45RO, CD127, CD197, TCRgd are specific to T memory cells, and the training was done without them. Overall, our model achieved better cell-wise and distributional accuracy assessed by Pearson correlation, R2 and EMD scores than cyMAE approach for all cell types (Table 2, Fig. S2). For T cell type, the performance was comparable.

**Table 2.**
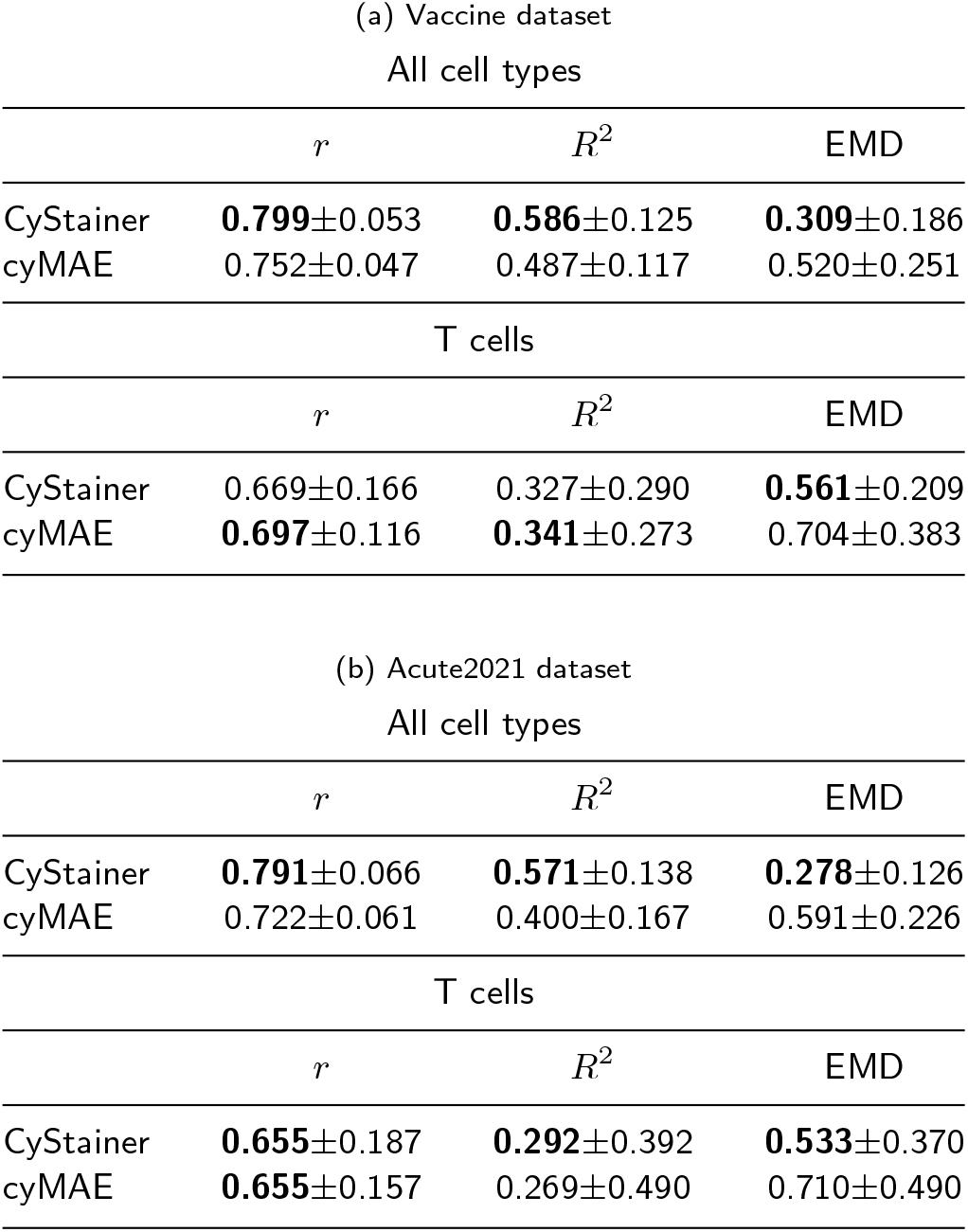
Comparison of marker prediction in unseen CyTOF COVID-19 data.

### Prediction in the absence of backbone markers

Finally, we asked whether the imputation can be achieved for a panel that does not share backbone with training datasets, as illustrated in Figure 4a. The test is designed based on the CytoBackBone data where backbone markers were omitted in panel E, while keeping the same panels A-D. In this test, we predicted all missing backbone markers for panel E at once. Here, only our transformer-based model could be evaluated due to incompatibility of model training scheme in other models. Specifically, to adapt cyCombine to this setup the training should be repeated including the previous and new data, while the input layer in cyMAE has a fixed marker configuration, limiting its application. Performance in this harder task still compared favorably to cyCombine imputation results for the leave-one-out task (r 0.522 vs 0.291; R-squared 0.233 vs -0.463) (Fig. 4b, Fig. S3, Table 1b).

**Figure 4.**
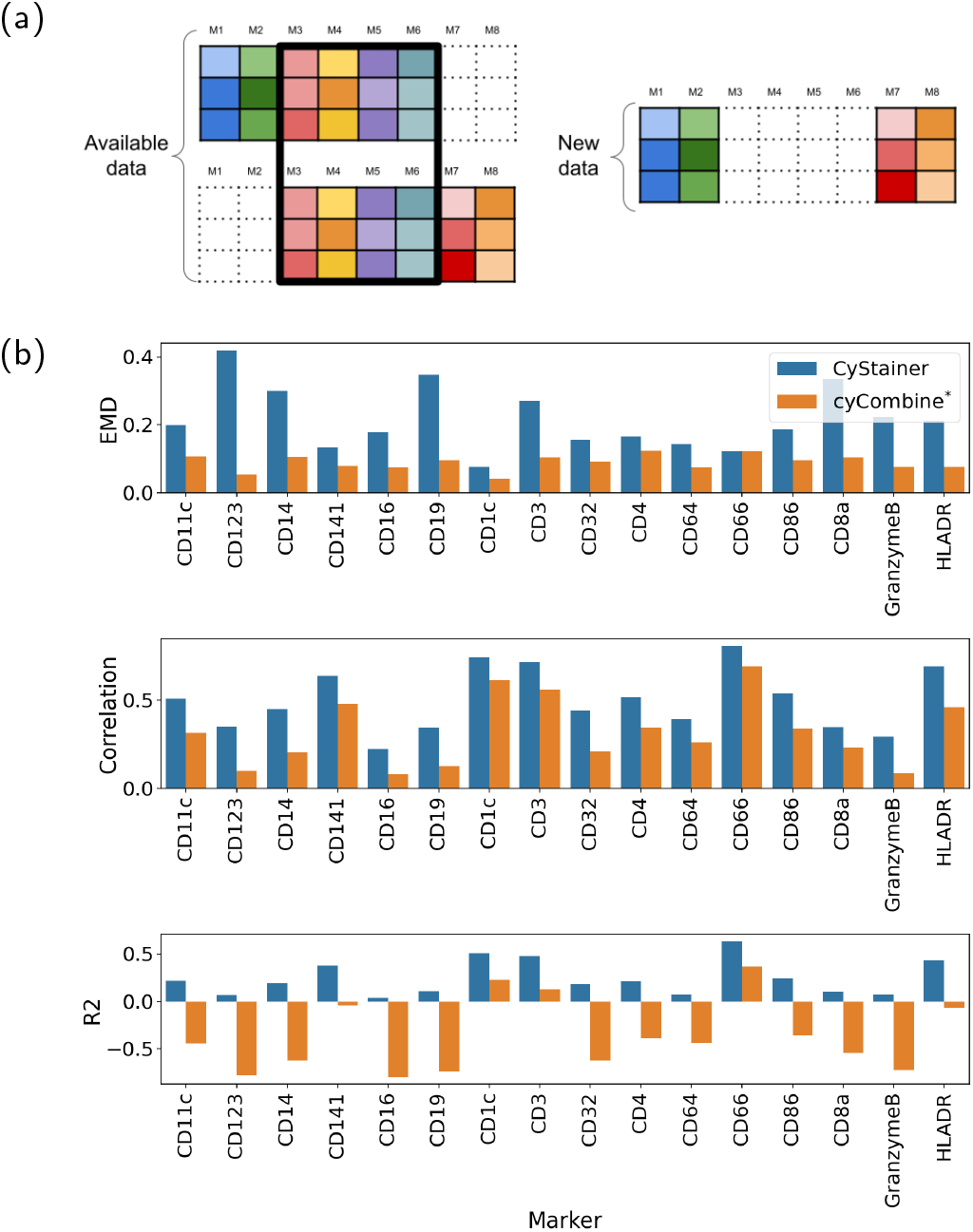
Performance comparison of CyStainer in backbone marker imputation. For reference, performance in leave-one-out marker imputation is shown for cyCombine^*\**^. (a) Backbone marker prediction scheme. (b) EMD, Pearson correlation and R-squared metrics marker wise.

### Evaluation of conditional accuracy

Next, we evaluated model performance in marker imputation that spans a larger set of markers and against additional models (GBR, cytoVI) using the bone marrow Infinity Flow dataset with 143 markers and 32 backbone markers. Consistently, CyStainer showed highest performance in cell-wise imputation accuracy, and ranked second in distributional accuracy (Table 1d). Of note, the strategy to predict markers in the original Infinity Flow study involved the training of separate GBR models for each marker, which is resource intensive, while our model as well as cytoVI can be trained in a straight-forward manner simultaneously for each marker.

While the overall performance gives a good indication of model performance, we chose to further examine the conditional accuracy for distribution matching, focusing on how well the marker profile is preserved within specific cell sub-populations. Since no labels were provided with the dataset, the different bone marrow cell populations were identified based on clustering (Supplementary fig. S4). Informative markers for each cluster were defined based on statistical comparison of each cluster to the rest. The EMD metric was then calculated for top five highly variable cluster markers, exemplified in Fig. 5. Overall, cyCombine exhibited the best distribution matching, followed by GBR, CyStainer, and cytoVI, with respective median EMD values of 0.073, 0.076, 0.087, and 0.118. Interestingly, well predicted markers-cluster combinations as well as poor predicted markers-cluster combinations mostly are consistent among different approaches.

**Figure 5.**
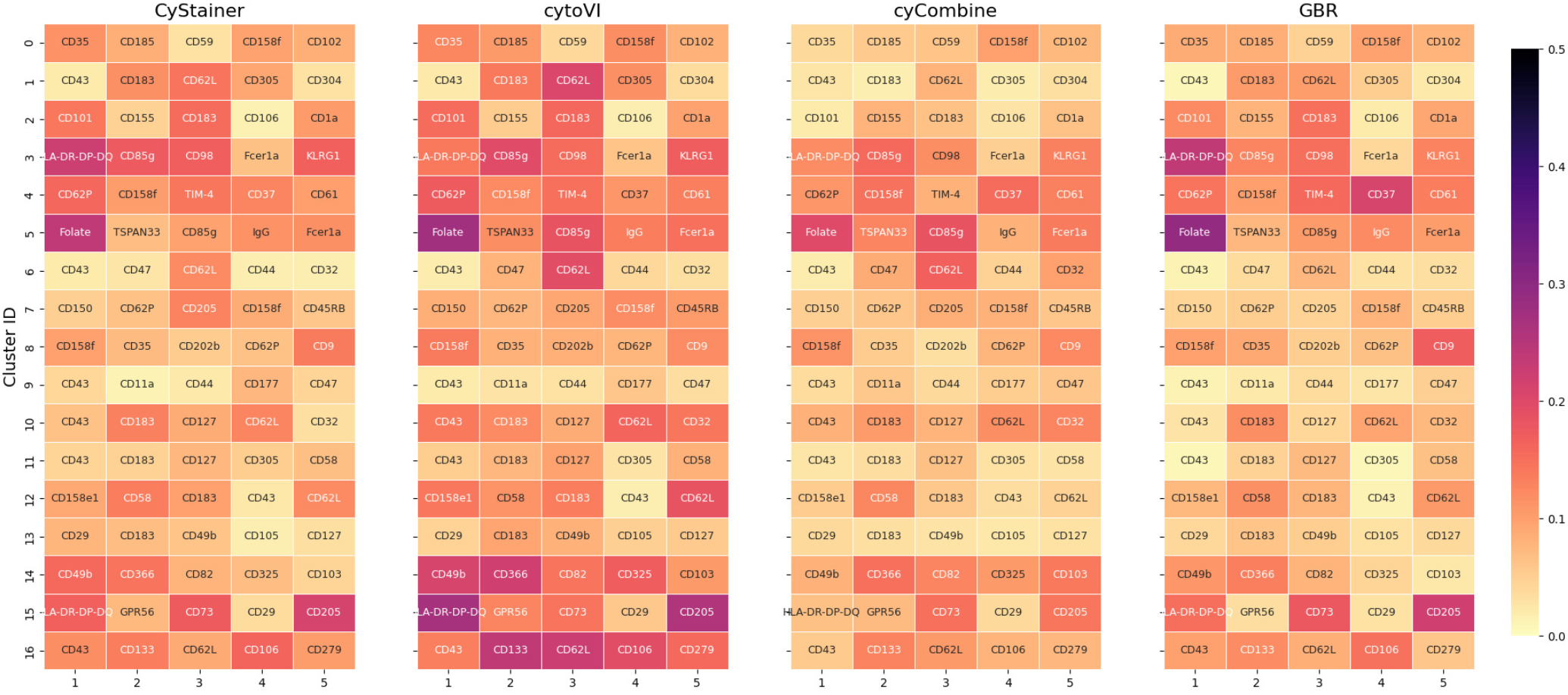
Conditional distributional accuracy comparing CyStainer, cytoVI, cyCombine, and GBR. The heatmap displays the EMD between the true and imputed distributions for the top 5 differentially expressed markers within each respective Leiden cluster.

### Evaluation of biological accuracy

Ultimately, to address biological or medical questions, it is important that model predictions preserve the biological interpretation. To benchmark model performance, we utilized the CITE-seq dataset from the NeurIPS 2021 multimodal data integration challenge including data generated at four different sites. We dropped one specific group of markers (Table S1) from each of the four sites. This dataset inherently contains also a significant batch effect, challenging the capacity of the models in handling additional technical variations. We then tasked CyStainer and cytoVI with predicting these missing markers specifically for each site. The cell- and distribution-wise imputation metrics are shown in Table 1e. In this dataset, CyStainer showed a clear advantage in distribution accuracy, registering the most optimal EMD scores across every marker group. It further demonstrated best cell-wise predictive accuracy in the majority of subsets, particularly within T cells (r 0.743 vs 0.735; R-squared 0.484 vs 0.398). This demonstrates CyStainer’s flexibility and capacity to perform reliable, site-specific marker imputation, even in the presence of technical variations.

To assess whether the imputed data preserved the separation of cell types, comparably to cell type labelling in the original study, we clustered the data using Leiden algorithm with resolution from 0.1 to 1.5 with step 0.1 (Supplementary fig. S5). For clustering we used the original measured markers defined for different immune cell types (Table S1) and compared this clustering to the labels and the clusters obtained with imputed values, as illustrated using Sankey plots for the B cell markers (Fig 6). The performance metrics are shown in Table 3. Based on ARI and NMI metrics our model prediction are comparable to the original values and outperforming cytoVI predictions.

**Table 3.** Comparison of clustering metrics across cell lineages in healthy bone marrow CITE-seq dataset. Average of ARI and NMI across all leiden clustering resolution is used. Best results are in bold, and second-best results are underlined.

| Model | T cell lineage |  | B cell lineage |  | NK and ILC |  | Monocytes/Mac/DC |  |
| --- | --- | --- | --- | --- | --- | --- | --- | --- |
|  | ARI | NMI | ARI | NMI | ARI | NMI | ARI | NMI |
| CyStainer | 0.393±0.129 | **0.626±0.069** | 0.109±0.111 | 0.353±0.108 | **0.089±0.092** | **0.220±0.077** | 0.148±0.103 | 0.437±0.132 |
| cytoVI | 0.345±0.134 | 0.538±0.076 | <u>0.123±0.122</u> | 0.327±0.097 | 0.058±0.032 | <u>0.172±0.049</u> | <u>0.174±0.126</u> | <u>0.456±0.142</u> |
| Original | **0.523±0.103** | <u>0.610±0.052</u> | **0.238±0.212** | **0.418±0.139** | <u>0.087±0.064</u> | 0.144±0.065 | **0.224±0.119** | **0.513±0.103** |

**Figure 6.**
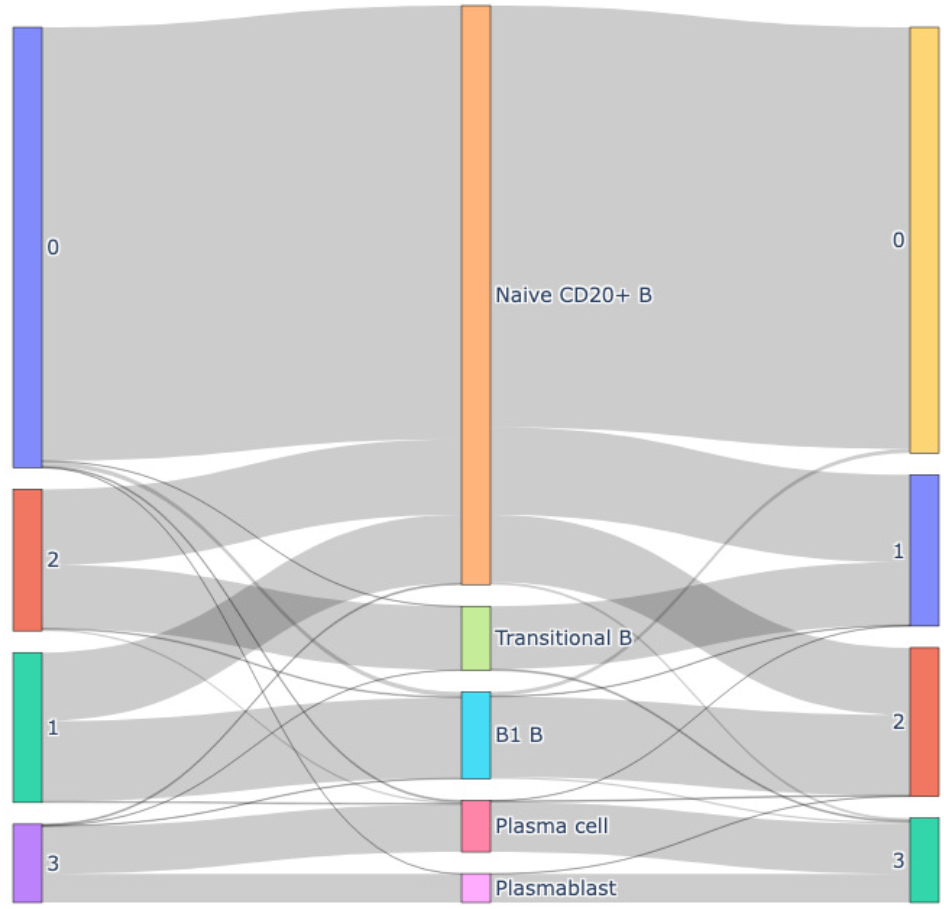
Sankey plot comparing the clustering based on B cell markers (left: original marker profile, right: predicted marker profile) compared to the annotated cell types in the Neurips CITE-seq dataset. The best Leiden clustering resolution (0.1) is defined based on ARI and NMI metrics.

Finally, to evaluate whether CyStainer can adapt to a new batch without suffering from catastrophic forgetting, we used two donors from site 1, 2 and 3 for training and one donor from each for testing, reserving site 4 data for validation. As in previous experiments, specific marker groups were dropped from sites 1, 2, and 3. The model was initially trained on the training data and evaluated on the test data. Subsequently, the model was fine-tuned 3 times on site 4, each time with one group of markers omitted. The prediction accuracy in test samples was evaluated for the missing markers both before and after fine-tuning to evaluate model resistance to catastrophic forgetting. The predictions for test samples before and after fine-tuning were almost exactly the same (Table S2). Furthermore, the prediction quality on site 4 after fine-tuning was comparable to that in the initial test samples, demonstrating CyStainer’s ability to successfully transfer learned knowledge across datasets with significant batch effect.

Taken together, the benchmarks carried out indicate that CyStainer has comparable accuracy (and often outperformed) existing solutions across multiple metrics. Our model design considerably simplifies the prediction workflow compared with multistage approaches proposed previously. The key novelty of our method is its ability to impute markers when only panels with no overlapping markers are available in training and test data, while high performance in preserving the biological information highlights its potential for biomedical applications to virtually stain missing sub-population markers.

## Discussion

The rapid expansion of high-parameter cytometry technologies has generated vast amounts of single-cell data. However, the lack of standardization across panel designs often constrained by clinical requirements or instrument limitations has created a significant bottleneck for data integration and comprehensive analysis. In this study, we introduced CyStainer, a novel transformer-based variational autoencoder designed to address these challenges through robust marker prediction.

The most significant limitation of existing marker imputation frameworks, such as cyCombine, cytoVI, cyMAE and GBR, is their reliance on a defined set of shared backbone markers or specialized experimental designs. Our results demonstrate that CyStainer successfully bypasses this requirement. By leveraging a transformer encoder to capture co-expression patterns, CyStainer can accurately impute markers even when training and test data lack overlapping panels. This capability represents a critical practical advancement, as it allows researchers to harmonize legacy datasets and varied clinical panels without needing to re-design their experimental setups.

Across multiple benchmark scenarios encompassing flow cytometry, CyTOF, CITE-seq, and Infinity Flow datasets, CyStainer consistently demonstrated competitive or superior performance compared to existing state-of-the-art models. Notably, CyStainer simplifies the computational workflow. While approaches like GBR require the resource intensive training of separate models for each individual marker, our architecture predicts all missing markers simultaneously. Furthermore, our targeted fine-tuning strategy enables the model to generalize effectively to unseen data subject to technical variation. By dynamically expanding the batch vocabulary while freezing the core network weights, CyStainer learns new batch effects without risking catastrophic forgetting of the global marker relationships established during initial training. This was successfully demonstrated by virtually staining unseen NeurIPS CITE-seq samples.

While CyStainer consistently ranked highest in cell-wise imputation accuracy (Pearson correlation and R-squared), we noted that the distributional accuracy assessed using EMD occasionally ranked slightly lower or second to other methods. Because EMD measures the overall distribution match rather than single-cell predictive accuracy, this suggests that while CyStainer is highly precise at predicting marker values for specific cells, there may be slight smoothing effects in the broader population distribution.

The flexibility and accuracy of CyStainer pave the way for the development of large-scale, pre-trained cytometry foundation models. Future iterations could be trained on massive, multi-institutional atlases, providing an off-the-shelf “virtual stainer” for clinical environments where panel sizes are heavily restricted.

## Data and code availability

ALL flow cytometry dataset is available under https://flowrepository.org/id/FR-FCM-Z68U.

Infinity Flow dataset is available under https://zenodo.org/records/15065910.

CytoBackBone CyTOF dataset is available under https://flowrepository.org/id/FR-FCM-ZYV2.

COVID-19 dataset is available under https://discover.pennsieve.io/datasets/401/version/1

NeurIPS CITE-seq dataset is available under https://www.ncbi.nlm.nih.gov/geo/query/acc.cgi?acc=GSE194122

The source package code is available under: https://github.com/sysgen-uef/cystainer_package.

Reproducibility code is available under: https://github.com/sysgen-uef/cystainer_reproducibility.

## Author contributions statement

K.I., V.H., and M.H. conceptualized the project. K.I., M.A.M., and S.R. curated the data. K.I. and M.A.M. conducted the formal analysis. K.I. and F.K. developed the methodology and software. K.I. produced the visualizations. F.K. and L.O. validated the results. O.L. provided the study resources. O.L. and M.H. acquired the funding. O.L., L.O., S.M., V.H., and M.H. supervised the research. K.I., V.H., and M.H. wrote the original manuscript draft. K.I., M.A.M., F.K., L.O., S.M., V.H., and M.H. reviewed and edited the manuscript.

## Acknowledgments

We thank Zhang Xuan and Salomonis Nathan authors of “An immunophenotype-coupled transcriptomic atlas of human hematopoietic progenitors” study for sharing raw infinity flow data upon our request.

This work was carried out with the support of UEF Bioinformatics Center, Biocenter Kuopio, Biocenter Finland, University of Eastern Finland, Finland.

