## Supplemental files for "CyStainer: A transformer-based variational autoencoder for robust marker imputation in high-parameter cytometry"

### Supplementary

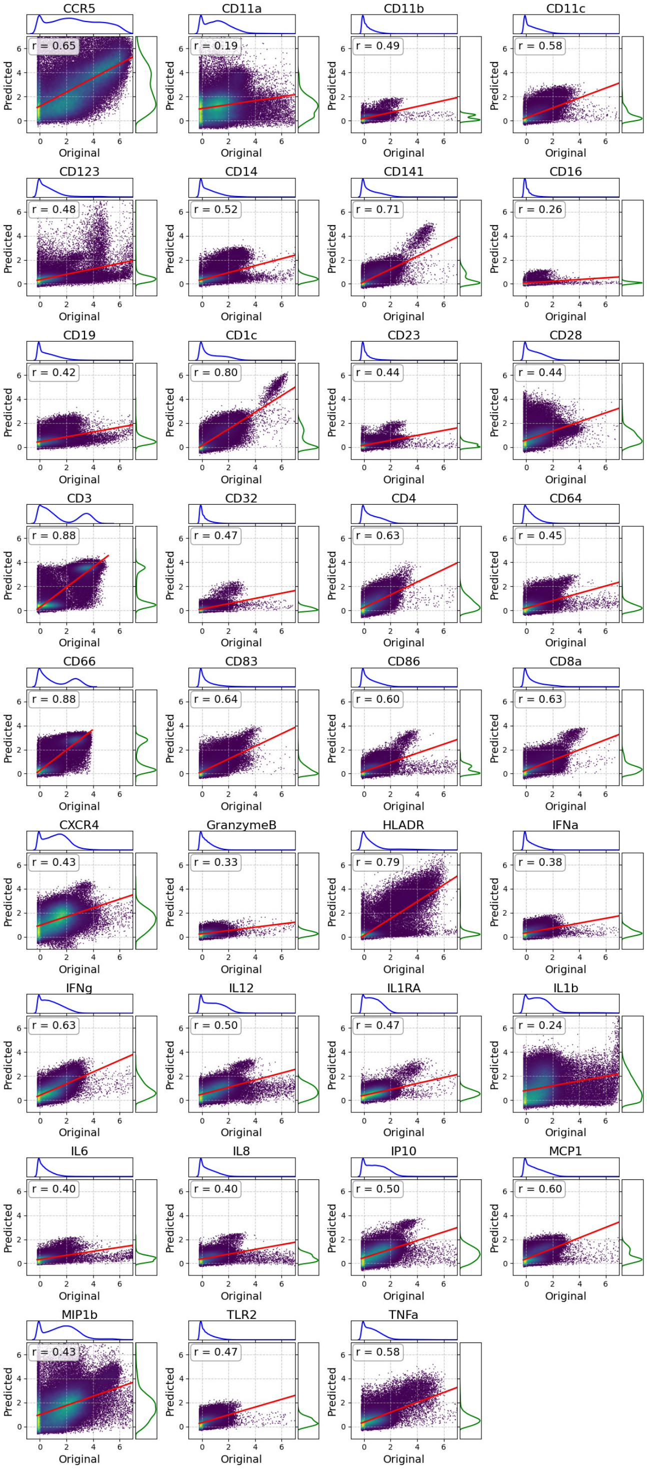


**Figure S1** Regplots of real versus CyStainer predicted values with Pearson correlation values for each marker in leave-one-out test for healthy PBMC CyTOF dataset

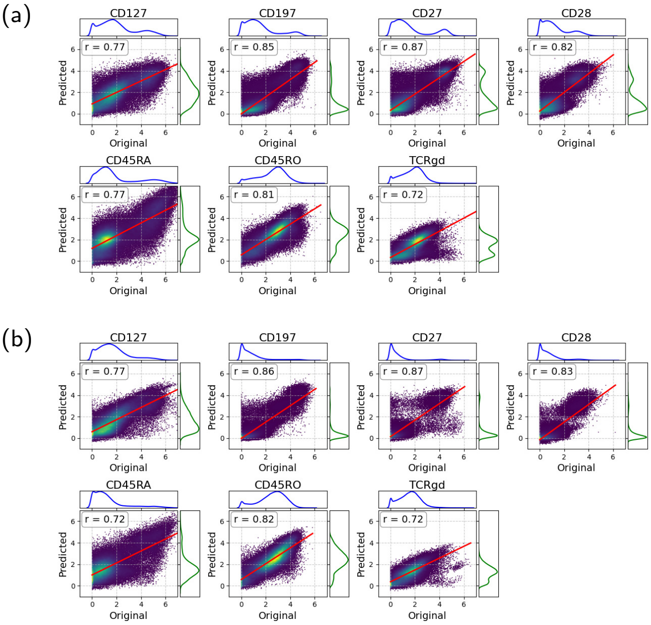


**Figure S2** Regplots of real versus CyStainer predicted values with Pearson correlation values for T cell markers for COVID CyTOF: (a) Vaccine dataset, (b) Acute 2021 dataset

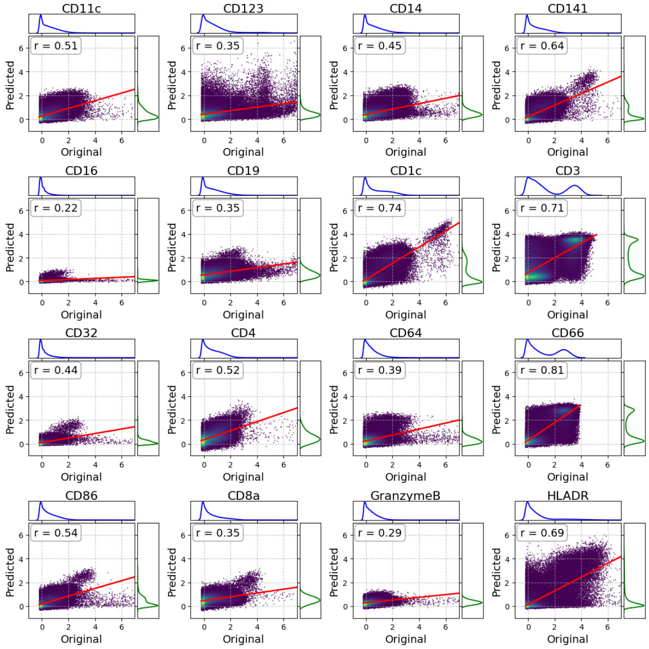


**Figure S3** Regplots of real versus CyStainer predicted values with Pearson correlation values for each marker in backbone prediction test for healthy PBMC CyTOF dataset

**Table S1.** Cell groups and corresponding markers

| Site | Cells | Markers |
|---|---|---|
| Site 1 | T-cell Lineage | CD3, CD4, CD8, TCR, TCRVa7.2, TCRVd2, CD2, CD5, CD7, CD25, CD27, CD28, CD127, CD134, CD137, CD154, CD223, CD278, CD279 |
| Site 2 | B-cell Lineage | CD19, CD20, CD21, CD22, CD23, CD24, CD72, CD79b, IgD, IgM, CD38, CD1c, CD1d, CD9, CD40, CD81, CD82, CD185, CD268 |
| Site 3 | NK and ILC | CD56, CD57, CD16, CD94, CD161, CD158, CD158b, CD158e1, CD314, CD335, CD85j, CD107a, CD122, CD226, CD244, CD272, CD352, KLRG1, TIGIT |
| Site 4 | Monocytes, Macrophages, DC | CD14, CD13, CD33, CD64, CD163, CD169, CD115, CD11b, CD11c, CD141, CD303, CD304, HLA-DR, CD32, CD83, CD86, CD88, CD172a, CX3CR1 |

**Table S2.** Healthy bone marrow CITE-seq fine-tuning.

| T cell markers | | | |
|---|---|---|---|
| | $r$ | $R^2$ | EMD |
| Site 1-3 trained | 0.742±0.236 | 0.501±0.332 | 0.061±0.032 |
| Site 1-3 fine-tuned | 0.742±0.236 | 0.501±0.332 | 0.061±0.032 |
| Site 4 fine-tuned | 0.645±0.253 | -1.147±4.184 | 0.079±0.064 |
| **B cell markers** | | | |
| | $r$ | $R^2$ | EMD |
| Site 1-3 trained | 0.665±0.169 | -0.070±1.367 | 0.047±0.027 |
| Site 1-3 fine-tuned | 0.665±0.169 | -0.070±1.368 | 0.047±0.027 |
| Site 4 fine-tuned | 0.774±0.135 | 0.240±0.714 | 0.055±0.040 |
| **NK and ILC cell markers** | | | |
| | $r$ | $R^2$ | EMD |
| Site 1-3 trained | 0.446±0.196 | -0.814±1.378 | 0.075±0.047 |
| Site 1-3 fine-tuned | 0.446±0.196 | -0.815±1.378 | 0.075±0.047 |
| Site 4 fine-tuned | 0.663±0.203 | -0.231±1.426 | 0.059±0.053 |

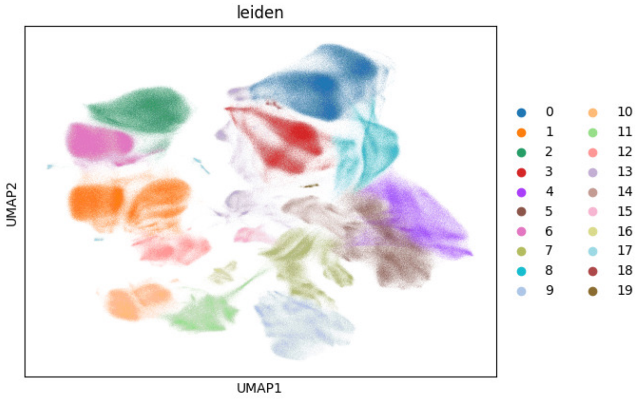


**Figure S4** UMAP projection colored by Leiden clustering for Infinity Flow dataset

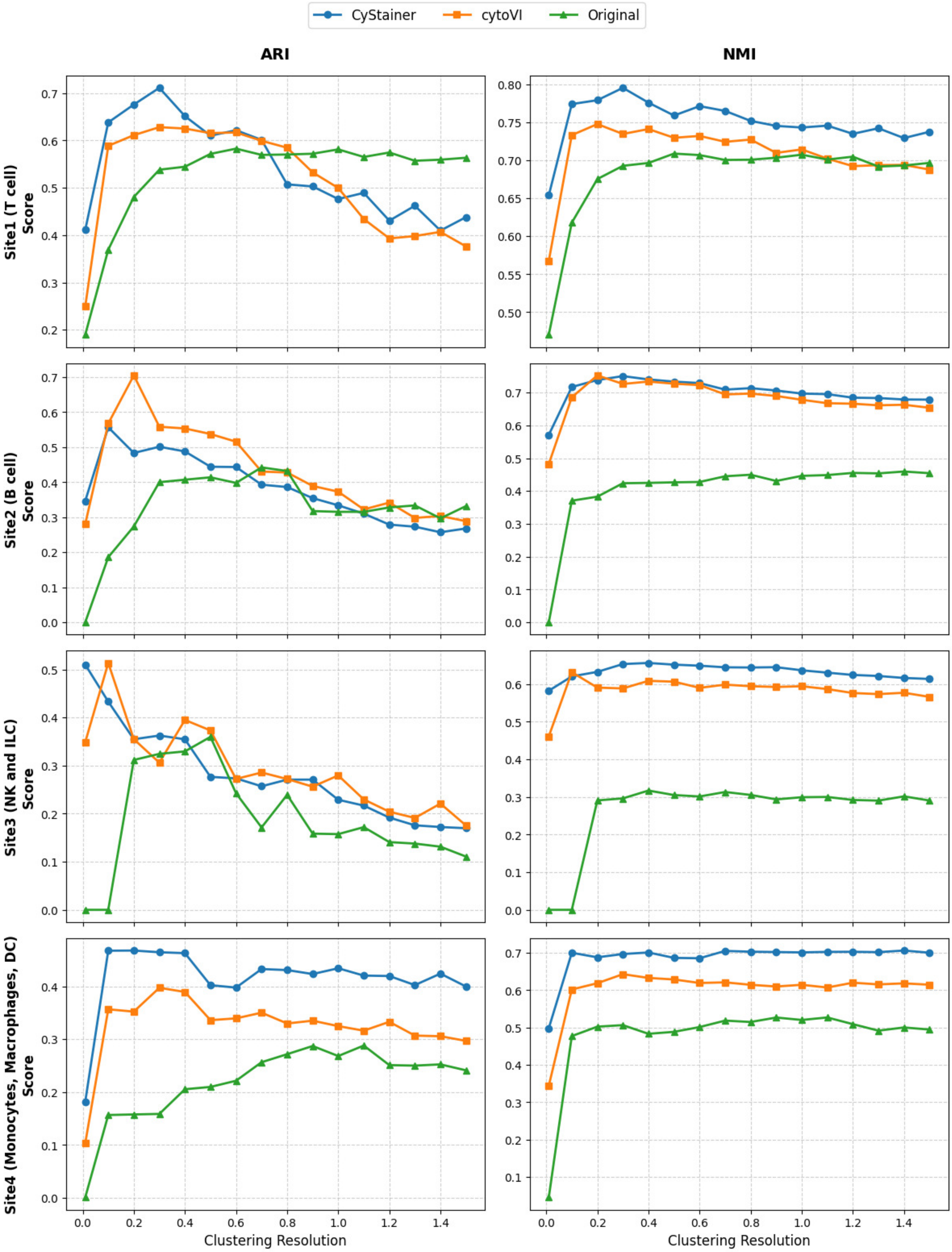


**Figure S5** ARI and NMI metric values for each Leiden clustering resolution. The clustering with asterisks is selected for Sankey Fig 6
